# Computational neural network provides naturalistic solution for recovery of finger dexterity after stroke

**DOI:** 10.1101/2021.06.22.449412

**Authors:** Ashraf Kadry, Sumner L. Norman, Jing Xu, Deborah Solomonow-Avnon, Firas Mawase

## Abstract

Finger dexterity is a fundamental movement skill of humans and the ability to individuate fingers imparts high motor flexibility. Disruption of dexterity due to brain injury reduces quality of life. Thus, understanding the neurological mechanisms responsible for recovery is critical to effective neurorehabilitation. Two neuronal pathways have been proposed to play crucial roles in finger individuation: the *corticospinal* tract, originating from primary motor cortex and premotor areas, and the subcortical *reticulospinal* tract, originating from the reticular formation in the brainstem. Finger individuation in patients with lesions to these pathways may recover. However, it remains an open question how the cortical-reticular network reorganizes and contributes to this recovery following a stroke. We hypothesized that interactive connections between cortical and subcortical neurons reflect dynamics appropriate for generating outgoing commands for finger movement. To test this hypothesis, we developed an Artificial Neural Network (ANN) representing a premotor planning input layer, a cortical layer including excitatory and inhibitory neurons and, a reticular layer that control motoneurons eliciting unilateral flexion of two fingers. The ANN was trained to reproduce “normal” activity of finger individuation and strength. Analysis of the trained ANN revealed that the natural dynamical solution was a near-linear relationship between the force of the instructed and uninstructed finger, resembling individuation patterns in humans. A simulated stroke lesion was then applied to the ANN and the resulting finger dexterity was assessed at multiple stages post stroke. Analysis revealed: (1) increased unintended force produced by uninstructed fingers (i.e., enslaving) and (2) weakening of the force in the instructed finger immediately after stroke, (3) improved finger control during recovery that typically occurs early after stroke, and (4) association of this behavior with increased neural plasticity of the residual neurons, as reflected by strengthening of connectivity weights between premotor and focal cortical excitatory and inhibitory neurons, but reduction in connectivity in shared cortical neurons. Interestingly, the network solution predicted that the reticulospinal pathway also contributed to the improved behavior. Lastly, the ANN also predicts the effect of cortical lesion size on finger individuation. Our model provides a framework by which to understand a number of experimental findings. The model solution suggests that a key mechanism of finger individuation is establishment of an interactive relationship between cortical and subcortical regions, appropriate to produce desired finger movement.

## Introduction

Humans, like other higher mammals, exhibit incredible finger dexterity. The skillful ability to move one or more fingers independently enables a large motor repertoire in mammals with prehensile digits. We rely on the ability to individuate fingers in a variety of daily activities, such as typing, tying shoelaces, or handling of utensils and various tools. Thus, any injury or pathology that interferes with finger individuation negatively impacts quality of life. After stroke, most people suffer from distal movement impairment which frequently manifests itself as both a decrease in finger strength and prominent deficit in hand dexterity. These are reflected by increased finger enslaving, or unintended force produced by the uninstructed fingers (i.e., inadequate finger individuation)^1–7^. Although a stroke patient may functionally recover the ability to flex and extend all fingers simultaneously, the finger individuation ability remains deficient. A longitudinal study that tracked finger individuation in stroke patients throughout the acute, sub-acute, and chronic phases revealed that recovery of finger individuation remains far from the level of healthy individuals and asymptotes after the first 3 to 6 months after the stroke event^5^. Nevertheless, the neural mechanism by which recovery of finger individuation occurs is still unclear.

Previous studies have shown the crucial role of the corticospinal tract (CST), originating from the primary motor cortex (M1) and the premotor cortex, in fractionated finger movements^8–10^. Experimental lesions or injury of the motor cortex or CST produce a serious detriment to individuated finger movements, resembling that seen in humans after stroke^8,11,12^. Recovery of finger dexterity from cortical lesions also reveals interesting results about the involvement of subcortical regions. In particular, the reticulospinal tract (RST), which originates from the reticular formation in the brainstem, was reported to undergo functional changes by modulating its activity after CST lesions during a fine independent finger movement task^13–15^ . Direct lesion of the brainstem medial RST, on the other hand, affected mainly posture, strength, and gross movements, while hand function remained unaffected^16^.

These observations suggest the existence of a neural circuit with interactive dynamics between the CST and RST that receives inputs (i.e., force and/or individuation commands) and produces the necessary motor output (i.e., individuated finger movement). Critically, changes in the connectivity at all levels seems to play a pivotal role in shaping recovery of finger movement after a brain lesion^15^. How exactly the cortical-reticular circuit reorganizes and contributes to the recovery process of finger individuation after stroke remains an open question.

We investigate how the cortical-reticular circuit reorganizes by developing a physiologically based computational model that is able to predict healthy behavior of finger movement, as well as behavior during the recovery period early after stroke. To date, most related works in modeling motor recovery have been limited to simulating either only wrist flexion force, or single-finger strength and individuation compared to the rest of the fingers^17,18^. In particular, Norman et al., 2017 presented a computational neural network model based on a stochastic reinforcement algorithm for a one-finger task and separately simulated the force patterns of the instructed finger (index) and the uninstructed finger (middle) in a non-lesioned normal mode, then independently in a lesioned stroke mode. Simulating the normal condition separately from the stroke condition limits the mechanistic understanding of how the network reorganizes and contributes to the recovery process of finger individuation after stroke. Here we present a complete solution that stems from a single simulation of a network at different conditions. This advance is crucial to better understand the possible neurophysiological mechanisms that might underlie stroke recovery.

In the present study, we built a novel Artificial Neural Network (ANN) model of the hand upper neuromotor system that is intended to simultaneously model two fingers, alternating between instructed and uninstructed modes with different force levels. Our ANN model captures residual capacity and dynamics at the cortical, subcortical and behavioral levels of finger recovery following a stroke. Importantly, our solution is complete in that once initialized to “normal” condition (i.e., prior to stroke), it is capable of simulating the different stages of the cortical motoneurons throughout the stroke event and the recovery process.

Notably, our ANN was trained to reproduce our proxy for the descending motor commands in a normal finger individuation condition, rather than the empirical behavioral and neuronal responses after stroke. We followed the normal anatomical connectivity constraints to impose physiologically-based structure on known features of cortical and subcortical connectivity^19,20^. This allowed the ANN to seek an optimum over a very broad range of dynamics, not limited by prior knowledge of finger recovery. The model findings also predict that post-stroke CST/RST integrity is correlated with improved finger dexterity recovery; a finding which can be validated in a clinical setting and, if successful, could inform patient treatment.

## Methods

### 1. Model Description

We developed a clustered ANN model constructed from 3 layers: input, hidden and output, (see ***Figure 1***, ANN Architecture Diagram, the architecture of the proposed ANN as implemented in our model). The input layer represents the commands for finger movements generated by motor cortical areas. The hidden layer, the computational heart of the model, represents the cortical primary motor neurons and the brainstem reticular neurons in the medulla/pons. The output layer represents the task action outcome generated by spinal motoneuron pools and muscles.

**Figure 1.**
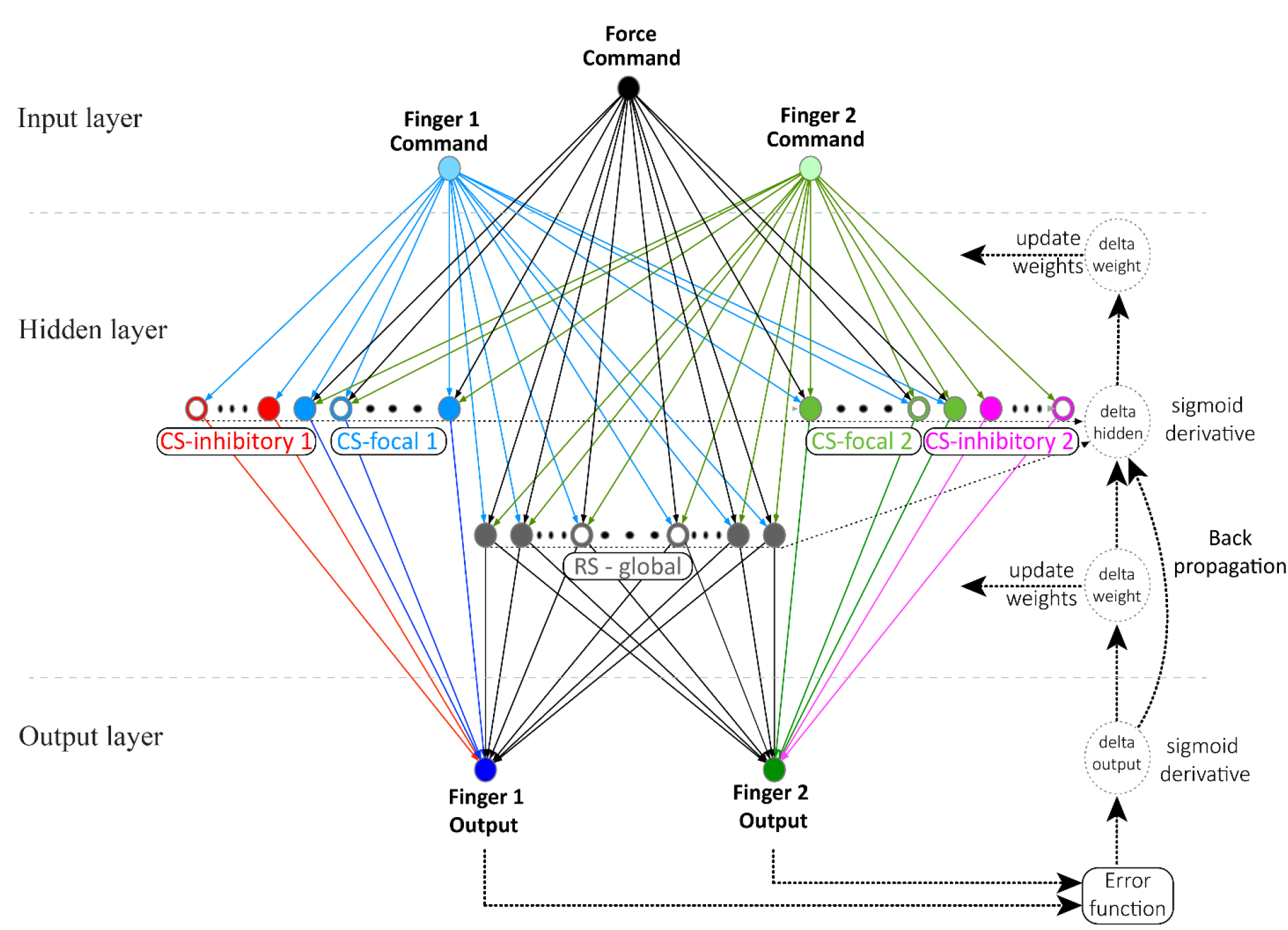
ANN Architecture Diagram. The inputs represent the pre-commands generated in the pre-motor cortex division relative to the two fingers. The hidden layers represent the primary motor cortex division (CS) and other sub-cortical motor regions (RS) involved in the control of voluntary movement. The outputs represent the outcome of the flexor and extensor neurons for each finger. Different colors represent different functions and/or a different finger. The neurons are represented by small circles (empty circles = neurons “disabled” by stroke, filled circles = healthy neurons). Bigger dotted circles and lines represent the back propagation flow. Abbreviations: CMD – command, CS – corticospinal, RS – reticulospinal.

#### 1.1. Input Layer

The input layer was composed of two types of inputs, two inputs for movement commands and one for force (see variables [*1*] and [*2*], Section 0). Each command input was fitted to a different finger, and the force represented the applied force level in the selected instructed finger. Four command movement tasks can be defined for the two fingers: instructed/uninstructed (1, −1), uninstructed/instructed (−1, 1), instructed/instructed (1, 1) and uninstructed/uninstructed (−1, −1). In this study, we focused on the first two commands (i.e., 1/−1 or −1/1) as they demonstrate individuation between the two fingers.

#### 1.2. Hidden Layer

The hidden layer was based on a simplified structure of the motor control areas, representing separable motor control functions and organized into different group types, four excitatory and inhibitory neuron groups for focal movement and one reticulospinal (RS) neuron group for gross movement. Each finger’s neuron cluster was a combination of focality groups and the RS group, which contained different excitatory and inhibitory neuron groups but shared the same RS group. The hidden layer state variables (see equation [*8*], Section 2.1) hold the intermediate hidden layer neurons’ output values.

#### 1.3. Output Layer

The output layer had two force outputs, each associated with a particular finger (see equation [*3*]). The outcome output was the combination of the instructed finger with the applied force per command.

#### 1.4. Input Layer to Hidden Layer Connectivity

The fingers’ command inputs were connected to all hidden layer neurons, with the exception that each command affiliated to one finger did not connect to the inhibitory neuron group of the other finger, and the force input was connected to the excitatory focal neuron groups and the shared RS neuron group. The strength of the connectivity between the layers was described by the connection weights (see equation [*6*]). A focal neuron in our network was defined as a neuron that could be driven by multiple neurons but is able to drive only one downstream neuron.

#### 1.5. Hidden Layer to Output Layer Connectivity

The hidden layer neurons were connected to the output layer neurons (outputs). Each finger was associated to one output, and the connectivity was based on the fingers’ cluster. Each finger output was driven independently by its cluster of neurons. Therefore, the focal neuron groups each drove their affiliated finger output, while the RS neurons group drove both outputs. Again, connectivity strength between the layers was represented by the connection weights (see equation [*9*]).

#### 1.6. Bias Parameters

Bias in Neural Networks is a mathematical operation that can be thought of as analogous to the role of a constant in a linear function, whereby the line is effectively translated by the constant value. Both the hidden layer and output layer neurons connect to bias constants. We added these constants to the sum of the inputs to the neurons and used them to shift the input values so that the outputs of the computation functions would fit within the desired range of output values. The bias is required when the summed weighted inputs of each neuron require adjustment before applying the activation function and helps the NN model to optimally fit data (see additional details in Section 0, model definition, in particular equations [*11*] and [*12*]).

### 2. Model Definition

The ANN model was characterized by three key features. 1) The number of residual motor cortex neurons in the hidden layers is inversely proportional to the magnitude of the overlap between the lesion and the motor area of the brain. When a stroke causes a large lesion in the motor cortex and/or CST, fewer cells contribute to the recovery process. 2) The force that each finger muscle generated was determined by the weighted sum activities of cortical and sub-cortical neurons in the hidden layers. Muscle force production is typically proportional to the firing rate of neurons in the motor cortex^17,21^. We therefore assumed that increase in the firing rate of a single neuron caused proportional increase in muscle force, up to a saturation limit, with the proportionality constant determined by the connection weights. 3) Lastly, we assumed that the motor system must find this solution by evaluating the results of task performance based on the deviation of the net force output of the fingers from the desired force targets (i.e., error function as teaching signal).

This type of reinforcement learning uses summary feedback of motor performance to update synaptic weights and can be achieved with computations implemented locally at synapses and is thus considered biologically plausible. We performed feed-forward-propagation followed by a back-propagation iterations algorithm to optimize for the results convergence. We compared the task performance in each feed-forward pass. and the error function was minimized during the back-propagation pass. The iterations were repeated until reaching or approaching the global minimum of the error function.

#### 2.1. Mathematical Definition

The command input variable, [*1*] *CMD_i,i_*=1,2, for each finger is used to capture a binary-type (instructed/uninstructed) command movement task that is required from each finger. The command information is encoded as +1/−1 rather than 1/0 to represent instructed/uninstructed movement tasks (a value of 0 does not work well with the ANN inputs, as all computation results would be zero regardless of its weights). The force input variable, [*2*] (*CMD*_3_ = *FRC*), is used as a percentage of full force and is provided as a number in the range of (0,1] (a value of 0 is not allowed, as movement cannot be achieved with zero force, a value of 1 represents 100% of the force). The force input is associated with the selected instructed finger based on the command inputs. The outputs of the ANN, [*3*] *FO_k_* = *f*([10])_*k*=1,2_ (see equation [*5*]), decode the task result representing the percentage value of force applied by each finger. The instructed finger will show actual instructed force, while the uninstructed finger will co-activate and show the uninstructed (i.e., involuntary) force. The expected outputs are derived from the command [*1*] and the force [*2*] and yield: [*4*] *EO_k_* = *FRC* × *CMD*_*k,k*=1,2_. We use the sigmoid, [*5*] 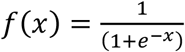, as the activation function. The sigmoid input values range is (−∞,+∞), representing the summed firing rates, and the output values, ranging in (0,1), represent the neuron’s firing rate percentage.

The command movement and instructed force inputs are weighted by the weight links connecting the inputs to the hidden layer neurons, [*6*] *WH*_*ij,i*=1,..,3;*j*=1,..*N*_. The weighted command movement and instructed force, [*7*] (*CMD_i_* × *WH_ij_*)_*i*=1,..3;*j*=1,..,*N*_, transforms to values in the range of [−1,1] and (0,1] respectively. These weighted inputs, [*7*] can be theoretically summed to values in the range of (−∞, +∞) feeding the corresponding activation function of each hidden layer neuron, [*5*]. This operation transforms the summed values back to values in the range (0,1) and represents the weighted activity of the neurons, hidden output [*8*] *HO_j_* = *f*([7])_*j*=1,..,*N*_. These intermediate outputs of the hidden layer neurons, [*8*], are saved as a vector of variables for later usage. Similarly, the intermediate outputs, [*8*], are weighted by the weight links connecting the hidden layer to the output layer, [*9*] *WO*_*jk,j*=1,..*N*;*k*=1,2_, and transformed to new values in the range (0,1), [*10*] (*HO*_*j*_ × *WO*_*jk*_)_*j*=1,..*N*;*k*=1,2_, and the sum values in the range (−∞, +∞) feeding the corresponding activation function of the output layer neurons; naturally the sigmoid function, [*5*], is used again to transform the summed values back to values in the range (0,1), and represent the output data, [*3*], (instructed/uninstructed finger) as a percentage of relative force of the movement (0 – no force, 1 – full force).

A bias is used to feed the hidden and output layer neurons, weighted using [*11*] *BH*_*j,j*=_1_,..,*N*_ and [*12*] *BO*_*k,k*=1,2_ weights, respectively. The weighted biases shift the neurons’ sigmoid activation functions, [*5*], to the desired range of values (see Section 1.6, Bias Parameters). Because the input to hidden layer weights, [*6*], must be positive values (reflecting excitatory activity), without bias the outcome values of the activation function, [*5*], range will then be limited to [0.5,1] (sigmoid of positive values). The hidden layer activation function, [*8*], was tuned to the range of (0,1] using the biased inputs, [*13*] [*BH*_*j*_+(*CMD_i_* × *WH_ij_*)_*i*=1,..,3_]_*j*=1,..*N*_, and equation [*8*] becomes: [*14*] *HO_j_* = *f*([13])_*j*=1,..,*N*_. The lacking range (0,0.5) can be reached only when applying the sigmoid to negative input values. Without bias, the weights start to switch to negative values (contradicting the excitatory context). Similarly for the output layer, despite the existence of the inhibitory neurons that enforce negative values, the summed inputs of the activation functions corresponding to these outputs still required tuning to the range (0,1], using [*15*] [*BO*_*k*_+(*HO*_*j*_ × *WO*_*jk*_)_*j*=1,..,*N*_]_*k*=1,2_, and equation [*3*] becomes: [*16*] *FO*_*k*_= *f*([15])_*k*=1,2_. There were very few inhibitory neurons (5% of the total focal neurons split between the two fingers).

In addition, we defined mask variables to control the interactions between different layers. They enforce connectivity limitations between different groups of neurons, as they represent different motor functions (simplified motor divisions) that do not necessarily directly interact. To model the interaction between the input and hidden layer groups of neurons, inhibitory mask, [*17*] *IM*_*ij,i*=1,...,3;*j*=1,..,*N*_, was defined (1^st^ masking vector (*i*=1) for first command, 2^nd^ masking vector (*i*=2 for second command and 3^rd^ masking vector (*i*=3) for the third command). Inhibitory neurons of one finger are affected by their associated finger command, but not by the other finger’s command. The 3^rd^ inhibitory mask vector ([*17*]; *i*=3) prevents the force [*2*] input from connecting to both inhibitory neuron groups. For the interaction between the hidden layer and finger output layer, finger mask, [*18*] *FM*_*jk,j*=1...*N*;*k*=1,2_, was defined. Focal neurons (excitatory and inhibitory) for each finger output are selected using its corresponding masking vector ([*18*]; *k*=*1,2*). The inhibitory neurons were negated using a dedicated hidden layer status variable, [*19*] *NS*_*j,j*=1,.,*N*_. A value of “1” indicates an excitatory neuron, while “−1” indicates an inhibitory neuron. A lesion was applied using the same status variable, [*19*], used for the hidden layer neurons. That is, each neuron was initialized to 1 or − 1 for healthy/active neurons and switched to 0 for “dead” (inactive) neurons due to stroke.

The error function, [*20*] *E_k_*= –(*E0_k_* – *F0_k_*)_*k*=1,2_, is defined as the difference between the expected output [*4*] and actual output [*16*] values for each finger separately. With back-propagation iterations using a gradient descent technique, we search for the optimal weights and minimize the error function [*20*]. We first calculate the derivative for the output layer outputs for the sigmoid function, [*16*], getting: [*21*] *∂O*_*k*_ = (*FO*_*k*_ × (1 – *F0*_*k*_))_*k*=1,2_, and the output delta error is: [*22*] *δO*_*k*_= (*E*_*k*_ × *∂O_k_*)_*k*=1,2_. Then we calculate backwards the weights of the connections between the hidden and output layers [*9*] using the delta correction: [*23*] *δWO*_*jk*_ = (*δO*_*k*_ × *HO*_*j*_)_*j*=1,..,*N*;*k*=1,2_. The output delta error, [*22*], is also backward propagated to the derivative of the hidden layer neurons sigmoid function, [*14*], getting: [*24*] *∂HO*_*j*_ = (*HO_j_* × (1 – *HO_j_*))*j*=1,..*N*, and the delta error correction is: [*25*] 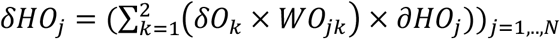. And finally, the inputs to hidden weights delta correction, [*26*] *δWH*_*ij*_ = (*δHO_j_* × *CMD_i_*)_*i*=1,...,3;*j*=1,..*N*_, is calculated and applied to update [*6*].

The individuation between the two fingers was defined as the ratio of the difference to the sum of the instructed (*FO*_1_) and uninstructed forces (*FO*_2_), [*27*] *I* = (*FO*_1_ – *FO*_2_)/(*FO*_1_ +*FO*_2_)^17^.

#### 2.2. Initialization and Parameter Setting

We set the global parameters to configure the main structure of the NN, e.g., number of hidden layer neurons (N=400), number of inputs (NI=3), and number of outputs (NO=2). We defined the dependent parameters to configure the inner structure of the NN, e.g., focality of the network (focal=0.4, 40%), inhibitory neurons (inhibit=0.05, 5% of the focal), and the rest of the neurons are the RS. We used the training and simulation related parameters to control the learning process, error correction resolution steps (eta=0.01), training cycles as number of days (*nDays*=360), and training repetition in each training cycle as daily dosage ([50, …, 50, 200, …, 200, 50, …, 50, 0, …, 0] vector of dosage values per day), defined differently for every stage of the training.

For setting up the NN structure we used mask and status variables, [17][18][19], based on the functions’ connectivity between the layers and pre/post stroke status as described in Section 1 (Model Description and ***Figure 1***, ANN Architecture Diagram) and defined in Section 0 (Mathematical Definition).

The initial weights of the NN were normally distributed numbers generated from the open interval (0,1) (~*normal*(*μ* = 0.5, *σ*^2^ = 1/12)) and are then masked using the mask variables, [17][18], while the non-connected weights were eliminated (set to “0”).

The bias constants were also weighted (generated similarly to the NN weights) to provide additional differentiation among different neurons’ activation functions. They were added as additional parameter inputs to the neurons of the hidden and output layers. The bias was used to compensate for the nature of the input values from one layer to another and the dependencies between the excitatory and inhibitory clusters, and adjust the distribution of the summed values within the desired range of the activation function for a better fitting. The bias setting process requires several trial- and- error tests before selecting good values for the NN model best fit. The hidden layer bias was set to “6” and the output layer bias was set to “−1”.

#### 2.3. Training and Simulation Methods

For training the ANN, not to be confused with, and not meant to simulate human motor training, the command targets and instructed force of the two fingers (*CMD*_1_, *CMD*_2_, *FRC*, respectively) are fed as inputs to the ANN. The fingers’ performance (actual response of the two fingers) were the actual result, i.e., the output of the ANN (*FO*_1_, *FO*_2_) as calculated in a feed-forward propagation. The error function was calculated and minimized in every iteration of the training process using gradient descent in a back-propagation flow (see Section 0, Mathematical Definition). The training for the initial normal condition is achieved by applying a multi-day and recurrent dosage force-based motor tasks to the normal (i.e., healthy) “pre-stroke” ANN, starting from an initially randomized state and converging to the desired instructed finger commands behavior. The simulation data is collected throughout the training process for later post-processing and demonstration.

We simulated a stroke by disabling a portion of the neurons in the hidden layer of the trained ANN, in proportion to the severity of the stroke, using the *NS* variable, [19]. To emphasize this, the lesion state was the actual outcome of the injured trained model without further training. The force-based motor commands were simulated at the stroke condition and the fingers’ outcome values were collected. In conjunction to the stroke, we reduce the learning capability (*η*) in accordance with the lesion severity. As assumed, injured brain plasticity is affected, and hence motor learning ability might be reduced.

Immediately after the lesion, the recovery process of the residual ANN represented the recovery that typically occurs at the early post-stroke phase and may continue to the chronic phase. The ANN is trained following the same method applied during the initial stage, and the simulation data is collected as well, but with the stroke condition as a starting point.

Since we are using all variants of movement commands for training, no additional validation is required for testing the converged NN; however, the quality of the NN convergence is highly dependent on initial values of the weights and may need several trials to reach the optimal NN for our different simulation usages. Configuring different ANN setups is done by setting new values for the ANN global parameters. In addition, the training can be tuned with number of days of recurrent loops with configured dosage iterations and learning factor.

#### 2.4. Training and Simulation Flow

The following steps are required to prepare the ANN database for simulation:

a. Configure the parameters for the desired ANN.
b. Select force (0,1] and initialize for chosen ANN setup/database.

- Select force ≠ 0 for simulation only using already saved initialization database.
- Use “0” to randomly generate new weights for new initialized training (this will also apply a full force), the new ANN database is saved.
c. Train the model at baseline normal state (pre-stroke), the ANN simulation data is collected.
d. Apply the acute-phase stroke condition by manipulating the number of neurons in the hidden layer of the NN.

- Deleting/blocking the function of some pool of neurons mimics the effect of the stroke in the brain.
- Severity of lesion would be represented by size of reduction in number of neurons.
- Run simulation and collect data (no training is applied at this step).
e. Apply the recovery and collect simulation data.

- This stage becomes the initial state of the chronic phase.
- Additional training for the finger tasks movement can be applied representing additional neurorehabilitation at chronic phase.
f. Compare collected simulation results with existing clinical study quality behavior.
g. Repeat b-f with different force cases to collect data for enslaving vs. force.
h. Repeat b-g with different stroke cases, stroke severity correlates with the lesion size and therefore the number of neurons deleted, and thus we represent different stroke cases.

### 3. Statistical Analysis

Statistical comparison between synaptic weights of the residual neurons before and after stroke was conducted using a paired two-tailed *t-test*. Specifically, we compared how the different weights were conditioned when re-trained after stroke. Significance level for all tests was set at 0.05.

## Results

### Finger strength and individuation in normal, lesioned and recovered condition

The ANN model is first initialized to approximately 50% of full force and zero individuation, starting with randomly generated weights, and then trained for two fingers (index and middle) to a pre-stroke condition by applying the instructed/uninstructed and force commands alternately in the same training set. At the end of this initialization process, the ANN is fully capable of the trained motor functionality of the two fingers for the two commands. The simulation data is collected throughout the training convergence and demonstrates the model behavior of this stage. The simulation shows the enhancement in the strength of the instructed/uninstructed fingers and individuation between them (maximizing the instructed force and minimizing the uninstructed force), and concurrently for the two fingers (see ***Figure 2*A**: command #1 simulation, pre-stroke-day, and ***Figure 2*B**: command #2 simulation, pre-stroke-day). The instructed force reaches 95.79% (index) and 96.48% (middle) of the maximum force, while the uninstructed involuntary force reaches 9.04% (middle) and 8.41% (index) of the maximum force. The individuation was measured as 0.83 and 0.84, respectively.

**Figure 2.**
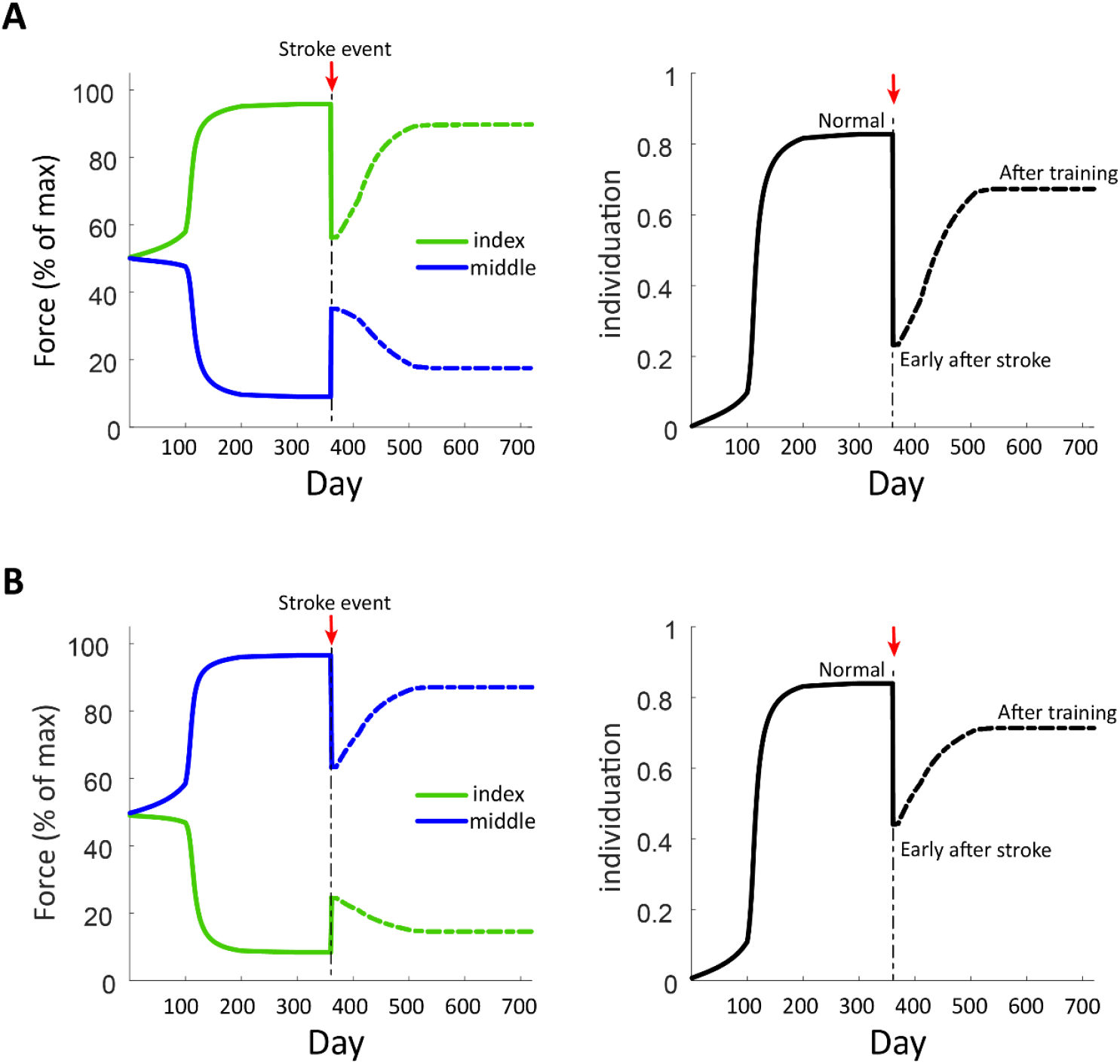
Fingers’ strength and individuation before and after stroke as predicted by the model. **A.** Command #1 Simulation (Index instructed, Middle uninstructed, Force 100%) simulating pre-stroke phase training, max instructed force and min uninstructed force (left), and full individuation (right), are achieved. Stroke event showing lesion acute phase degradation in instructed force and increase in uninstructed/unintentional force, leading to individuation reduction between the two fingers. The recovery early after stroke demonstrates enhancement in instructed and uninstructed force behavior and thus individuation recovery. **B.** Command #2 Simulation (Index uninstructed, Middle instructed, Force 100%) results and behavior are similar in qualitative manner to A.

A stroke event is applied by deleting 40% of the neural network from the hidden layers (see Section 0, Model Definition). The simulation with the two commands is executed at this point and exhibits the behavior of the impaired model via a drop in the instructed force, rise in the uninstructed force and detriment to the individuation between the two fingers (see ***Figure 2*A**: Command #1 Simulation, stroke-day, and ***Figure 2*B**: Command #2 Simulation, full force, stroke-day). The instructed force reaches 56.36% (index) and 63.37% (middle) of the maximum force, while the uninstructed force reaches 35.12% (middle) and 24.52% (index) of the maximum force. The individuation was measured as 0.23 and 0.44, respectively.

Following the stroke event, i.e., proceeding from the stroke condition state, the model is trained using the same training method to regain some enhancement of the motor behavior and to represent the recovery early after stroke. Similar to the pre-stroke stage, simulation data is captured throughout the training process up to the limit of the impaired ANN convergence. We observe a rise in the instructed force and drop in the uninstructed force, and eventually enhancement in the individuation (see ***Figure 2*A**: command #1 simulation, post-stroke recovery, and ***Figure 2*B**: command #2 simulation, post-stroke recovery). The instructed force reaches 89.65% (index) and 87.06% (middle) of the maximum force, while the uninstructed force reaches 17.51% (middle) and 14.55% (index) of the max force. The individuation was measured as 0.67 and 0.71, respectively.

### Increased co-activation of uninstructed finger as a function of instructed finger strength

Next, we sought to explore the relationship between the force of the uninstructed finger for different strength amplitudes of the instructed finger. To test this relationship in our model, we repeated the simulations with different force targets and measured the uninstructed involuntary force from the middle finger and the instructed force from the index finger in the different phases (normal, early after stroke during recovery training, and post-recovery training) of these simulations for each force target. In ***Figure 3***, we plotted the uninstructed vs. instructed forces (normalized to max force in our simulation, the clinical results were plotted as measured). The y axis represents the co-activation (i.e., involuntary forces) produced by the uninstructed finger in accordance with the applied force of the instructed finger as shown on the x axis. The slope ratio represents the individuation ability of the instructed finger. We see that early after stroke, the involuntary force of the uninstructed finger increases (i.e., reduced individuation) to more than it was in the normal/non-paretic case in both the model and stroke patient graphs.

**Figure 3.**
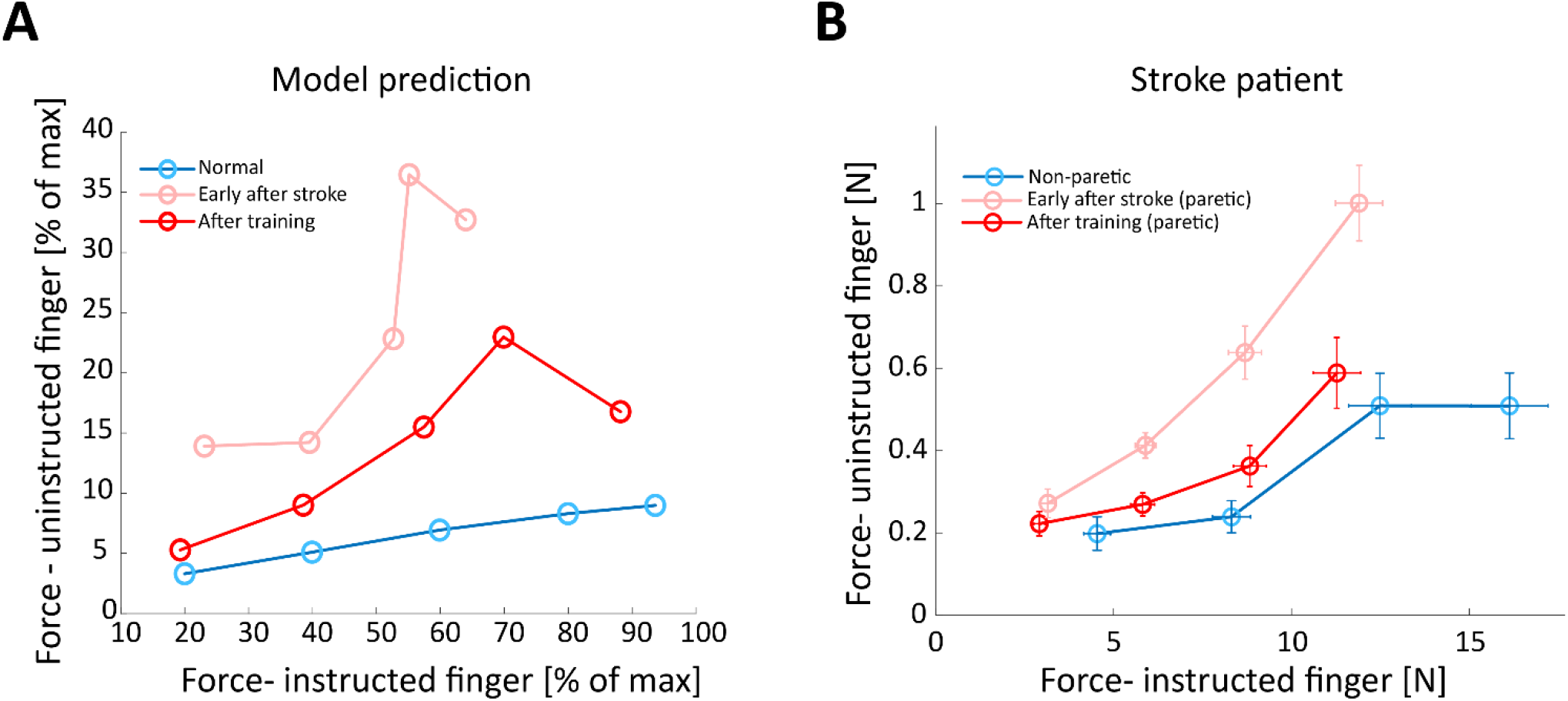
Uninstructed forces as a function of instructed finger strength. Model prediction (left) vs. clinical measurements (right): Recovery effect on finger individuation as predicted by the model and as observed in a stroke patient. **A.** Forces in the uninstructed finger plotted against the force generated by the instructed finger at multiple force amplitudes as predicted by the model. Colored lines represent normal pre-stroke phase (blue), acute phase (light red), post-recovery (induced spontaneously or with training early after stroke) and chronic phase (dark red). Model parameters: (Index instructed, Middle uninstructed, Stroke: 40%, Forces: 20%, 40%, 60%, 80%, 100%). **B.** Reduced enslaving in the individuation task in a stroke participant (data from Mawase et al., 2020). Forces of the non-instructed finger as a function of the forces in the instructed finger for the non-paretic hand reflecting the pre-stroke baseline level (blue), early after stroke (light red) and after training (dark red).

Interestingly, after recovery (e.g., induced by spontaneous recovery and/or additional rehabilitative training), we observed that the model predicted reduced involuntary uninstructed force (i.e., increased individuation) that got closer to a normal level (***Figure 3* A**, Model Prediction). Quantitively, this reduction was captured by the slope of a linear regression line that was fitted to each data set and showed an almost flat line (slope=0.078, with 95% CI of 0.06-0.09) in the normal condition (i.e., before stroke), substantial increase of slope (slope=0.55, with 95% CI of −0.09-1.192) immediately after stroke and significant reduction (slope=0.22, with 95% CI of −0.04-0.049) after recovery. This uninstructed force-finger strength relationship replicated what we have previously reported in human stroke patients (***Figure 3* B**, Stroke Patient: shows actual data from a stroke participant during an individuation task^7^).

### Effect of lesion size on finger individuation

We measured the effect of lesion size on finger individuation during the acute and sub-acute recovery phases. A large lesion in our model is presumed to reflect a large infarct in the cortical and/or CST regions. Our model predicted that very small lesion size (e.g., up to 10% of the total neurons in the hidden layer) results in almost no degradation in forces or individuation. With 20% lesion size, we see a small effect of the stroke in the acute phase, but the model is still capable of regaining almost all its motor capabilities after the recovery training process. Lesion sizes in the range of 30%-70% result in a decrease in finger individuation in the acute phase followed by improvements in motor function that asymptote to levels below full recovery. These below-normal-function levels are apparently related to the severity of the stroke and the amount of training induced by the model. Finally, the model predicted that lesion sizes in the range of 80-90% cause a severe drop in motor function that cannot be effectively restored, and additional training enhancements are very limited (***Figure 4*A**).

**Figure 4.**
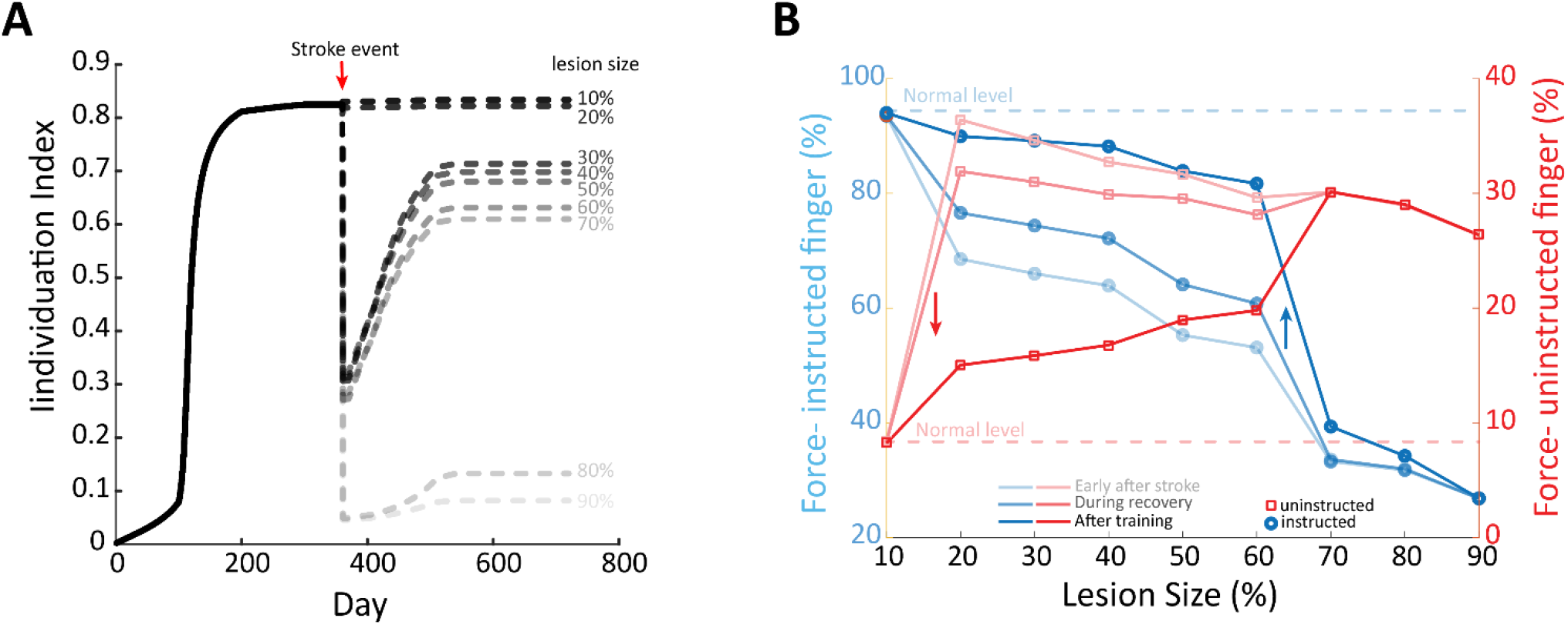
Effect of lesion size on instructed vs uninstructed fingers. Model parameters: (Index instructed, Middle uninstructed, Stroke 10-100%, Force 100%). Simulations: All are trained similarly during pre-stroke phase, then stroke with different lesion sizes is applied. We recorded the results early after stroke at sub-acute phase, then during recovery in chronic phase and at the end of rehabilitation effort. **A.** Individuation between Index and Middle fingers in accordance with lesion size. The graph demonstrates that the greater the lesion size, the more severe the stroke effect on individuation, and the less likely recovery is after stroke. Lesion sizes <20% have minimal effect on the individuation and recovery. **B**. The dynamics of fingers’ strength following lesion with different sizes. In the instructed finger (blue lines), the greater the lesion size, the less instructed force that can be produced by the model, starting with the light blue capturing the forces early after stroke, the darker blue during recovery and darkest blue at end of rehabilitation process. Recovery increases from the acute post-training phase, i.e., increase in instructed force. The uninstructed force (light red lines) is inversely affected by the lesion size early after stroke and during recovery. After training, however, the uninstructed force (dark red) is directly affected by lesion size, as seen by reduced uninstructed force levels compared to what is predicted early after stroke.

We tested a resulting assumption that effect of lesion size on finger individuation is driven by the relative changes between instructed and uninstructed fingers. To test this, we measured the effect of lesion size as predicted by the model on the instructed and uninstructed forces early after stroke, during the recovery process and after recovery completion. We plotted the uninstructed forces and the instructed forces vs. lesion size on the same plot for comparison (***Figure 4* B**). We see that the instructed force values are always inversely related to the lesion size, i.e., they have similar slope direction in the different measurement states, however, the slope is steeper early after stroke, less steep during recovery and becomes more moderate towards the end of training. The uninstructed force values, however, show a different behavior. While the plots early after stroke and during recovery have similar slope-direction compared to the instructed force plots slopes, indicating an inverse relationship to the lesion size. We found that after training, the slope was inversed, and the uninstructed force was directly relative to the lesion size. Overall, this suggests that the recovery of finger individuation is directly related to lesion size, signifying reduction in the capability to reduce the uninstructed force.

### Changes in synaptic weights in residual CST and RST during stroke recovery

The observation of improved finger individuation after stroke is proposed to be associated with plasticity changes in network connectivity of the residual neurons. Specifically, we investigated how the connectivity strength of the model changed during the recovery process, i.e., how the different weights were conditioned when re-trained after stroke. We evaluated a representative case, 100% force and 40% stroke, which demonstrates the recovery enhancements in both force and individuation of the two fingers (***Figure 5***).

We found increased plasticity of the residual neurons, as reflected by gain increase (paired *t-test*, *p* < 0.0001) in the command to the instructed finger’s focal weights (i.e., connectivity between premotor and focal cortical excitatory neurons, ***Figure 5* A**). Although it did not reach statistical significance (*p* = 0.18), the connectivity for the focal inhibitory neurons showed a similar trend of positive gain (***Figure 5* B**). On the other hand, we found reduction in connectivity in shared cortical neurons (*p* < 0.0001, ***Figure 5* B**). The network solution predicted that the reticulospinal pathway (i.e., RST) also contributed to the improved behavior (***Figure 5*D**). This is demonstrated by significant increase (*p* = 0.0181) of synaptic weight in the force input of the RS neurons. No change (*p* = 0.9) was detected in weights between the command and the RS neurons (***Figure 5* E**). ***Figure 5* F** summarizes the re-organization of the residual neurons of the network that contributed to the recovery of finger individuation after stroke. Improved finger individuation was achieved by strengthened connectivity of focal excitatory cortical neurons, weakened shared excitatory cortical neurons of the other finger and strengthened connectivity of RS neurons.

**Figure 5.**
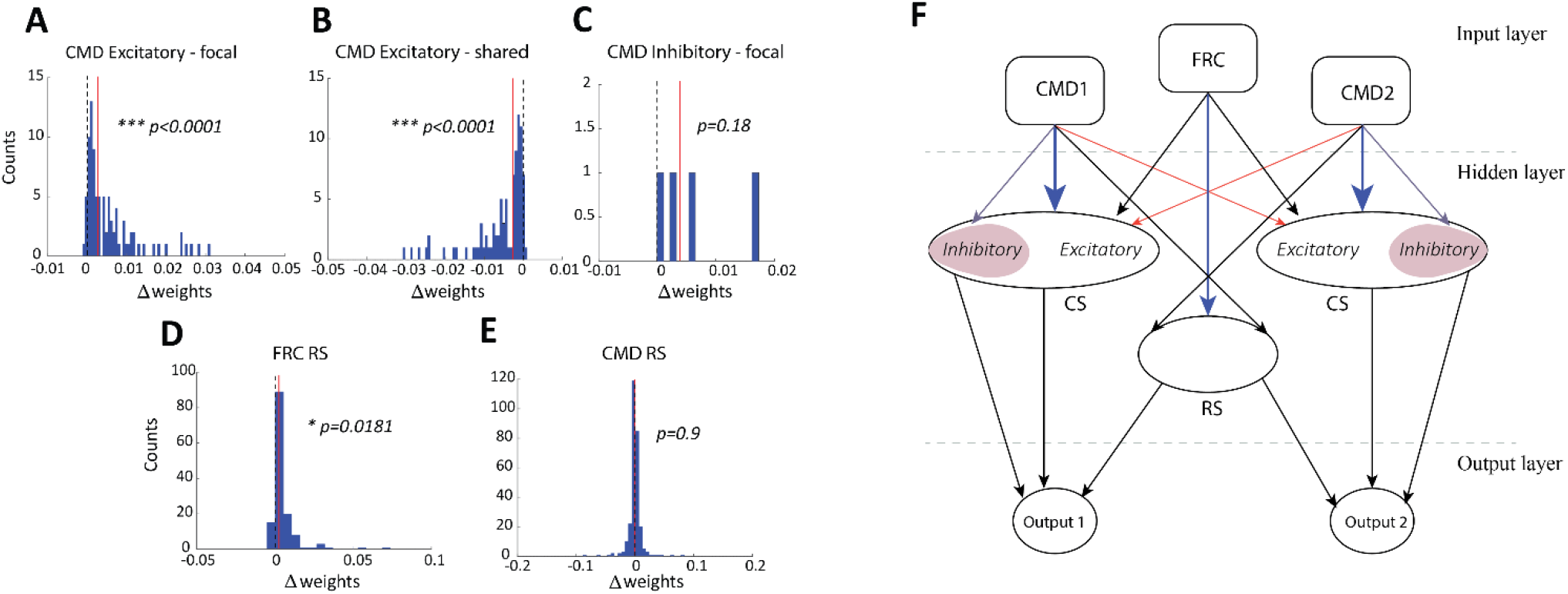
Weight Gain Histograms. Model Parameters: (Index instructed, Middle uninstructed, Stroke 40%, Force 100%). **A.** CMD to excitatory focal (direct-finger) weights gain increased. **B.** CMD to excitatory focal (opposite-finger) weights decreased. **C.** CMD to inhibitory focal weights gain increased. **D.** CMD to RS weights gain almost unchanged (slightly increased). **E.** FRC to RS weights gain increased. **F.** Re-organization of the impaired network during recovery of finger individuation after stroke. Blue arrows – increased connectivity, red arrows-reduced connectivity and black arrows-no significant change in connectivity. Abbreviations: CMD – command, FRC – force, RS – reticulospinal. * indicates p<0.05, *** indicates p<0.0001.

## Discussion

In the present study, we designed an ANN model with physiologically-based architecture that provides a naturalistic solution for recovery of finger dexterity after stroke. Our central result is that an ANN trained to produce finger individuation exhibited dynamics that strongly resemble that of healthy individuals and patients after having recovered from a stroke. The resemblance between the model outcome and reported data in previous clinical works was manifested by the substantial reduction of finger individuation immediately after stroke^2,7^, the recovery pattern following training, the near-linear relationship between uninstructed forces and instructed finger strength and the relationship between lesion size and severity of impairment in finger individuation^5,6,7^. Notably, this agreement was not achieved by fitting the ANN to actual clinical data. Rather, the agreement between model outcome and clinical data emerged as a result of the architecture of the excitatory/inhibitory cortical CS and subcortical RS neuron pools needed to generate the normal patterns of individuation. And as mentioned earlier, once initialized to pre-stroke condition, our solution is capable of simulating dynamic functional capacity of the cortical motoneurons throughout the lesion event and the recovery process that follows. In addition, the model makes predictions that might provide mechanistic explanation about the functional reorganization of the cortical and subcortical network during recovery of control of finger movement.

Our modeling study provides a framework by which to understand a number of experimental findings related to finger dexterity. First, the pattern of instructed forces, uninstructed forces and individuation in a normal condition, as seen in ***Figure 2***, mimics the convergence relation between the instructed to uninstructed force levels as observed in humans and primates^8,10^. Second, the immediate ANN response to the simulated stroke event revealed increased involuntary uninstructed forces that were driven by weakening of the instructed finger and exaggerated force of the uninstructed finger. This is in agreement with documented clinical observations of acute post-stroke phase finger functionality in which post-stroke patients exhibit reduced instructed finger force capacity and increased uninstructed forces^3,22^. Third, the model exhibited how the impaired motor system re-adjusted and learned new neural activation of the residual cells to compensate for the loss in finger control during the recovery process. Behaviorally, we observed an increase in the instructed force and decrease in the uninstructed force, and eventually enhancement in finger individuation during the recovery period. This is in line with recent studies that showed meaningful improved, yet incomplete, finger individuation during the early months after the stroke event^5,22^.

### Cortical-subcortical neural basis of finger individuation

Our model predicts post-lesion plasticity in both cortical and subcortical areas for recovery of finger dexterity. While strengthening of the connectivity in the residual descending cortical pathway seems to contribute to a larger extent to fractionating finger movement, strengthening of the reticulospinal tract seems to compensate for the loss of finger strength^15^. Inspection of changes in synaptic weights during stroke recovery as predicted by the model revealed plasticity in the residual CST and RST. Specifically, we observed strengthening of the weights to the focal cortical excitatory and inhibitory neurons, that together control the desired finger movement. Strengthening these weights indicates enhancing the individuation and some extent of force. On the other hand, we reported weakening of the weights that connected the focal neurons to the other finger. This reduced contribution inversely affected the individuation, and thus weakening them is in favor of individuation. Paradoxically, the model predicts that strengthening of the non-focal brainstem connectivity also constrains the individuation recovery due to the faciliatory effect of such increased connectivity on strength of the uninstructed finger. Specifically, we found that weights connecting to RS neurons that mainly serve for applying force are strengthened. The weights of the connection between force command and RS neurons apparently do not add to the generated force or individuation in these conditions. This striking feature of model prediction mirrors neurophysiological findings in previous primate models^15^.

Many previous studies in human and animal models provide evidence that the corticospinal tract and reticulospinal tract underlie finger dexterity and strength, respectively^24^. In 1968, Lawrence and Kuypers examined the effects of lesions to the CST and found that the capacity for independent movements of the hand digits was lost after a complete bilateral lesion of the pyramidal tract; while strength was severely impaired after bilateral disruption of the RST^8,16^. In human stroke, Xu and colleagues analyzed diffusion tensor imaging (DTI) data and measured finger individuation and strength and found that lesions in the hand areas in M1, as well as the CST, correlated more with impaired individuation than with strength in humans with stroke^5^. A recent follow-up TMS study demonstrated the reliance of finger individuation recovery on the integrity of CST as measured by the presence of motor evoked potentials in the hand^25^. Zaaimi et al., 2012^15^ found that connections from medial brainstem pathways (probably largely reticulospinal tract) undergo functional changes after corticospinal lesions, and that the connection strengthening was selectively specific for inputs to forearm flexors, but not extensors that were left unchanged. This asymmetric muscle-specific pattern of recovery has been widely seen in stroke patients^26,27^. After initial paralysis, stroke patients show increased activity of flexor muscles, including finger flexors, that sometimes developed into abnormal flexion synergies. Activity of the extensor muscles, in contrast, remains very weak and unchanged^1^. The (in)ability to voluntarily activate finger extensors is a reliable biomarker predicting functional outcomes^27^.

### Severity of impairment in finger dexterity correlated with lesion size in the motor cortex and motor-related subcortical areas

There is mounting evidence suggesting that lesion size within specific brain areas might be a major factor in the ability to restore motor function after stroke, and the improvement, or lack thereof, in motor activity^28–30^. Several studies have demonstrated that greater damage to the corticospinal projections is associated with more impairment in stroke patients^31–33^. Our model demonstrates the correlation between lesion size in motor areas and reduced plasticity of the injured brain, which explains the (in)ability to restore motor function in these cases. In our model, a very small lesion size around 10% does not induce impairment in the motor function. This might be explained by the fact that the NN model is converged to a “mathematically” stable global minimum and requires a substantial “hit” to be disturbed. This aspect of our model is in accordance with clinical data of stroke, as the effects on function of mild strokes can be difficult to quantitatively assess ^34–37^. In the range of 20% to 70% lesion size, we can observe the effect of the stroke event in our model. The modelled acute phase emphasizes the level of the disturbed motor function relative to the size of the injured brain region and shows that there is still room for a certain amount of spontaneous recovery of motor capabilities, but that it is limited inversely to the lesion severity. In our model, lesion size with more than 80% dead neurons demonstrates severe impairment of motor function that is almost incapable of being restored and minimally or not responsive to rehabilitation. The clinical analogue has been documented in the literature with poor functional outcomes for severe stroke and a more difficult challenge to design and implement rehabilitation protocols capable of inducing improvements in this population^38,39^. Nevertheless, neurophysiological quantification of residual CST neurons that survived after the stroke, as well as association between this quantity and motor impairments, require future research with high-resolution imagining tools.

Our model predicts a decrease in uninstructed force, as a function of lesion size, immediately upon stroke event and early after stroke (i.e., acute and sub-acute phases). A possible explanation for this is that the model’s post-stroke “motor system”, including CST and RST, has less available neurons that can affect force, and since the involuntary forces of the uninstructed finger are linearly affected by the amount of applied force in the instructed finger, when less instructed force can be generated by the model’s “motor system”, it will lead to less uninstructed force as well. When rehabilitative training is applied, the slope of the uninstructed force is inverted. This can be explained by the model’s ability to adjust the remainder of the intact neurons, thus allowing higher gain with smaller lesion sizes. In the case of small lesions (e.g., less than 60%), reduction in uninstructed force might be driven by changes in the residual inhibitory cortical neurons. Nevertheless, our model predicts that when the lesion size is large (e.g., above 60%), most of the inhibitory neurons are removed, leaving no room for enhancement, and thus limiting the reduction of the uninstructed force after training.

The individuation, as we observe in ***Figure 4* A**, is calculated based on the normalized difference between instructed and uninstructed forces. Immediately and early after stroke, the magnitude of the decrease in the instructed forces across different lesion sizes is larger than the magnitude of the increase in the uninstructed forces; therefore, individuation decreases as lesion size increases. Recovery induced by training and/or occurring spontaneously caused increase in the forces of the instructed finger and decrease in the uninstructed forces. This can be explained as enhancement in the focal segments, excitatory and inhibitory weights, with the excitatory neurons positively affecting the instructed forces and, conversely, the inhibitory neurons negatively affecting the uninstructed forces.

Altogether, these results indicate that relatively simple dynamics between cortical and subcortical neurons could provide a naturalistic explanation for the recovery of finger dexterity. It is essential to understand which subsystems contribute to recovery of finger movement in order to provide a rational basis to develop circuit-level therapeutic strategies that will optimize rehabilitation. Some aspects of the reported finding are in accordance with well-reported clinical and neurophysiological outcomes, but others provided mechanistic prediction of the interactive relationships in the neural network that underlie finger dexterity. These predictions can be tested in future research working primarily with human and/or animal models that typically exhibit finger dexterity.

### Limitations and future directions

Our model has some limitations. First, in this work we limited our model to two fingers (index and middle). In addition, the hidden layer segments of the NN, focal CS and RS neurons, were uniformly split between the two fingers’ function divisions and have similar connectivity between the different layers, and thus have the same capacity of capabilities and dependencies, i.e., the two fingers are mutually inclusive in achieving the instructed forces and enslaving each other. In addition, the stroke was applied to all segments of the NN neurons with equal weight (same percentage). These assumptions do not necessarily represent the real organization of the motor system, nor does it reflect how a real stroke lesion may differently affect motor divisions and representation of multiple fingers. In fact, it was shown that control of individuated finger movement is widely distributed in the primary motor cortex^2^, and electrical stimulation often elicited involuntary movements of multiple fingers^10,40^. Finally, our model is an oversimplification of the proposed network that potentially underlies recovery of finger dexterity. Simplification of the proposed ANN model was pronounced in its architecture, including design, connections, and training. Thus, although our model predicated plausible outcomes that highly resembled data from human and/or primate research, it seems that more complex architecture of neural networks including additional brain areas, beyond cortical and RST, must be involved in control of finger movement.

As for future directions, though the ANN model was limited to two fingers for our simulation requirements, the model is easily scalable to support all fingers and can be adjusted to support individual forces (different force levels) of the instructed fingers. In this case, the level of complexity of the model increases, and the training set must be revised accordingly, and thus simulation run time increases substantially. Such enhancements and/or adding further motor-related divisions may be optimally addressed using one of the commonly used advanced computations. More advanced neural network models or deep learning frameworks might be used and trained to simulate enhancements in both strength and individuation in hand motor function, based on existing clinical experiments and available data (e.g., size of lesion and affected part/s of the brain motor divisions).

Our model makes clinically-testable predictions. For example, it predicts that post-stroke CST/RST integrity is correlated with improved finger dexterity recovery. This prediction could be verified in a clinical study wherein CST/RST tractography from diffusion-weighted MRI is used to predict patient outcome. This protocol could validate this model finding and possibly lead to future clinical work that stratifies patients into therapeutic interventions based on CST/RST tractography and the model’s predictions of expected recovery. Thus, ultimately, treatment can be planned based on the desired target goals for finger individuation and/or strength. The improvement, or lack thereof, in the motor activity as predicted by the model will help us estimate the amount of motor recovery of the training dataset.

## References

1. Twitchell TE. 1951. The restoration of motor function following hemiplegia in man. Brain 74(4):443–480.

2. Schieber MH, Poliakov A V. 1998. Partial Inactivation of the Primary Motor Cortex Hand Area: Effects on Individuated Finger Movements. Journal of Neuroscience 18(21):9038–9054.

3. Li S, Latash ML, Yue GH, et al. 2003. The effects of stroke and age on finger interaction in multi-finger force production tasks. Clinical Neurophysiology 114(9):1646–1655.

4. Lang CE, Schieber MH. 2004. Reduced Muscle Selectivity during Individuated Finger Movements in Humans after Damage to the Motor Cortex or Corticospinal Tract. Journal of Neurophysiology 91(4):1722–1733.

5. Xu J, Ejaz N, Hertler B, et al. 2017. Separable systems for recovery of finger strength and control after stroke. Journal of Neurophysiology 118(2):1151–1163.

6. Wolbrecht ET, Rowe JB, Chan V, et al. 2018. Finger strength, individuation, and their interaction: Relationship to hand function and corticospinal tract injury after stroke. Clinical Neurophysiology 129(4):797–808.

7. Mawase F, Cherry-Allen K, Xu J, et al. 2020. Pushing the Rehabilitation Boundaries: Hand Motor Impairment Can Be Reduced in Chronic Stroke. Neurorehabilitation and Neural Repair 34(8):733–745.

8. Lawrence DG, Kuypers HGJM. 1968. The functional organization of the motor system in the monkey: I. The effects of bilateral pyramidal lesions. Brain 91(1):1–14.

9. Lemon R. 1988. The output map of the primate motor cortex. Trends in Neurosciences 11(11):501–506.

10. Schieber MH. 1990. How might the motor cortex individuate movements? Trends in Neurosciences 13(11):440–445.

11. Chapman CE, Wiesendanger M. 1982. Recovery of function following unilateral lesions of the bulbar pyramid in the monkey. Electroencephalography and Clinical Neurophysiology 53(4):374–387.

12. Schwartzman RJ. 1978. A Behavioral analysis of complete unilateral section of the pyramidal tract at the medullary level in macaca mulatta. Annals of Neurology 4(3):234–244.

13. Riddle CN, Edgley SA, Baker SN. 2009. Direct and indirect connections with upper limb motoneurons from the primate reticulospinal tract. Journal of Neuroscience 29(15):4993–4999.

14. Soteropoulos DS, Williams ER, Baker SN. 2012. Cells in the monkey ponto-medullary reticular formation modulate their activity with slow finger movements. The Journal of Physiology 590(16):4011–4027.

15. Zaaimi B, Edgley SA, Soteropoulos DS, Baker SN. 2012. Changes in descending motor pathway connectivity after corticospinal tract lesion in macaque monkey. Brain 135(7):2277–2289.

16. Lawrence DG, Kuypers HGJM. 1968. The functional organization of the motor system in the monkey: II. The effects of lesions of the descending brain-stem pathways. Brain 91(1):15–36.

17. Norman SL, Lobo-Prat J, Reinkensmeyer DJ. 2017. How do strength and coordination recovery interact after stroke? A computational model for informing robotic training. In: IEEE International Conference on Rehabilitation Robotics. p 181–186.

18. Norman SL, Wolpaw JR, Reinkensmeyer DJ. 2020. Targeting Neuroplasticity to Improve Motor Recovery after Stroke. bioRxiv :2020.09.09.284620.

19. Koeppen BM, Stanton BA. 2008. Berne & Levy Physiology, 6th ed. Elsevier.

20. Tortora GJ, Derrickson BH. 2010. Principles of Anatomy and Physiology, 13th ed. Wiley.

21. Ashe J. 1997. Force and the motor cortex. Behavioural Brain Research 86(1):1–15.

22. Jo HJ, Maenza C, Good DC, et al. 2016. Effects of unilateral stroke on multi-finger synergies and their feed-forward adjustments. Neuroscience 319:194–205.

23. Schieber M, Lang C, Reilly K, et al. 2009. Selective Activation of Human Finger Muscles after Stroke or Amputation. Advances in experimental medicine and biology 629:559.

24. Baker SN. 2011. The primate reticulospinal tract, hand function and functional recovery. Journal of Physiology 589(23):5603–5612.

25. Schambra HM, Xu J, Branscheidt M, et al. 2019. Differential Poststroke Motor Recovery in an Arm Versus Hand Muscle in the Absence of Motor Evoked Potentials. Neurorehabilitation and Neural Repair 33(7):568–580.

26. Cauraugh J, Light K, Kim S, et al. 2000. Chronic motor dysfunction after stroke: Recovering wrist and finger extension by electromyography-triggered neuromuscular stimulation. Stroke 31(6):1360–1364.

27. Fritz SL, Light KE, Patterson TS, et al. 2005. Active finger extension predicts outcomes after constraint-induced movement therapy for individuals with hemiparesis after stroke. Stroke 36(6):1172–1177.

28. Habegger S, Wiest R, Weder BJ, et al. 2018. Relating acute lesion loads to chronic outcome in ischemic stroke-an exploratory comparison of mismatch patterns and predictive modeling. Frontiers in Neurology 9(SEP):737.

29. Yoo AJ, Barak ER, Copen WA, et al. 2010. Combining acute diffusion-weighted imaging and mean transmit time lesion volumes with national institutes of health stroke scale score improves the prediction of acute stroke outcome. Stroke 41(8):1728–1735.

30. Vogt G, Laage R, Shuaib A, Schneider A. 2012. Initial lesion volume is an independent predictor of clinical stroke outcome at day 90: An analysis of the Virtual International Stroke Trials Archive (VISTA) database. Stroke 43(5):1266–1272.

31. Schaechter JD, Perdue KL, Wang R. 2008. Structural damage to the corticospinal tract correlates with bilateral sensorimotor cortex reorganization in stroke patients. NeuroImage 39(3):1370–1382.

32. Lindenberg R, Renga V, Zhu LL, et al. 2010. Structural integrity of corticospinal motor fibers predicts motor impairment in chronic stroke. Neurology 74(4):280–287.

33. Schulz R, Park C-H, Boudrias M-H, et al. 2012. Assessing the Integrity of Corticospinal Pathways From Primary and Secondary Cortical Motor Areas After Stroke. Stroke 43(8):2248–2251.

34. Cole M. 1993. Measurement in neurological rehabilitation. Neurology 43(2):459.

35. Carlsson GE, Möller A, Blomstrand C. 2003. Consequences of mild stroke in persons <75 years-A 1year follow-up. Cerebrovascular Diseases 16(4):383–388.

36. Duncan PW, Samsa GP, Weinberger M, et al. 1997. Health status of individuals with mild stroke. Stroke 28(4):740–745.

37. Edwards DF, Hahn M, Baum C, Dromerick AW. 2006. The Impact of Mild Stroke on Meaningful Activity and Life Satisfaction. Journal of Stroke and Cerebrovascular Diseases 15(4):151–157.

38. Nakibuuka J, Sajatovic M, Nankabirwa J, et al. 2015. Early mortality and functional outcome after acute stroke in Uganda: prospective study with 30 day follow-up. SpringerPlus 4(1):450.

39. Pereira S, Graham JR, Shahabaz A, et al. 2012. Rehabilitation of individuals with severe stroke: Synthesis of best evidence and challenges in implementation. Topics in Stroke Rehabilitation 19(2):122–131.

40. Tanji J, Okano K, Sato KC. 1988. Neuronal activity in cortical motor areas related to ipsilateral, contralateral, and bilateral digit movements of the monkey. Journal of Neurophysiology 60(1):325–343.

